# Auditory Attention Implements Complementary but Independent Cortical Mechanisms of Enhancement and Suppression

**DOI:** 10.64898/2026.04.27.720989

**Authors:** Max Schulz, Christopher Gundlach, Jonas Obleser, Malte Wöstmann

## Abstract

In noisy environments, goal-directed behavior requires the brain to prioritize task-relevant sound and to suppress distraction. While mechanisms of *proactive* attention in the presence of foreknowledge about targets and distractors are well-characterized, the neural dynamics governing *reactive* auditory attention in the absence of foreknowledge remain less understood. To fill this gap, we recorded the electroencephalogram (EEG) from *N* = 34 participants performing an auditory search task. Participants identified an amplitude-modulated spoken target number amid distractor numbers of varying salience. Presenting stimuli from three spatial locations enabled isolation of lateralized electrophysiological responses to targets and distractors. We quantified selection of targets versus distractors by continuously tracking participants’ cursor movements on a virtual response pad. Behaviorally, the target-directedness of cursor movements (i.e., *towardness)* increased for target repetitions (positive priming) and decreased when the target was a distractor from the previous trial (negative priming), indicating respective enhancement and suppression. Neurally, lateral targets evoked an N2ac component reflecting reactive enhancement. Importantly, we here demonstrate for the first time in an auditory attention task that distracting sounds evoke an auditory distractor positivity (P_D_) component. Target enhancement and distractor suppression were largely independent: their neural and behavioral markers were not significantly correlated and they originated from distinct cortical sources. Furthermore, shorter N2ac-but not P_D_-latencies related to higher towardness. By dissociating neural signatures and behavioral relevance of enhancement and suppression, our findings refine biased-competition models of attention, demonstrating that reactive selection relies on separable mechanisms rather than a single competitive gain process.

**Significance statement:** Everyday listening requires the rapid selection of relevant sounds and ignoring distraction, even when there is no prior knowledge of what will be heard. Here, we reveal the neural mechanisms that enable such reactive auditory attention. Combining electroencephalography with continuous measures of behavior, we show that target enhancement and distractor suppression are supported by distinct neural processes, indexed by the N2ac and the auditory P_D_ component. These processes are largely independent, with only enhancement predicting behavior. This dissociation challenges unitary accounts of attentional selection and instead supports models in which separable mechanisms of enhancement and suppression operate in parallel. Our findings provide a mechanistic framework for understanding how the human brain flexibly prioritizes information in complex listening environments.

## Introduction

Humans navigate complex environments where sensory inputs compete for limited cognitive resources. Since peripheral information exceeds the capacity of the central nervous system, goal-directed action requires enhancing task-relevant inputs and/or suppressing task-irrelevant distractions (Desimone & Duncan, 1995). Depending on available foreknowledge, these processes occur either proactively (pre-stimulus), or reactively (post-stimulus; Braver, 2012; Geng, 2014; Liesefeld et al., 2024). While the neural basis of reactive attention is well-characterized (Eimer, 2014; Liesefeld et al., 2017; Luck et al., 2021; Theeuwes, 2010), auditory attention models lack a mechanistic implementation of reactive enhancement and suppression.

A mechanistic understanding of attention requires differentiating attentional capture (increased processing; Theeuwes, 2025) and suppression (decreased processing; Gaspelin & Luck, 2018; Lui et al., 2023, 2025). These processes correspond to distinct temporal neural response profiles (illustrated in Fig. 1 A). While target enhancement manifests as a neural response increase (Andersen & Müller, 2010; Eimer, 1996), distractor processing can distinguish competing theoretical models: (1) initial capture followed by disengagement without suppression, (2) a cascade of early capture followed by suppression, or (3) suppression without preceding capture. Evidence suggests enhancement and suppression are independent in vision (Khodayari et al., 2026) and proactive auditory attention (Wöstmann et al., 2019). This study quantifies reactive auditory enhancement and suppression to determine whether distractor suppression is functionally independent or coupled with target enhancement (Fig. 1B).

**Figure 1.**
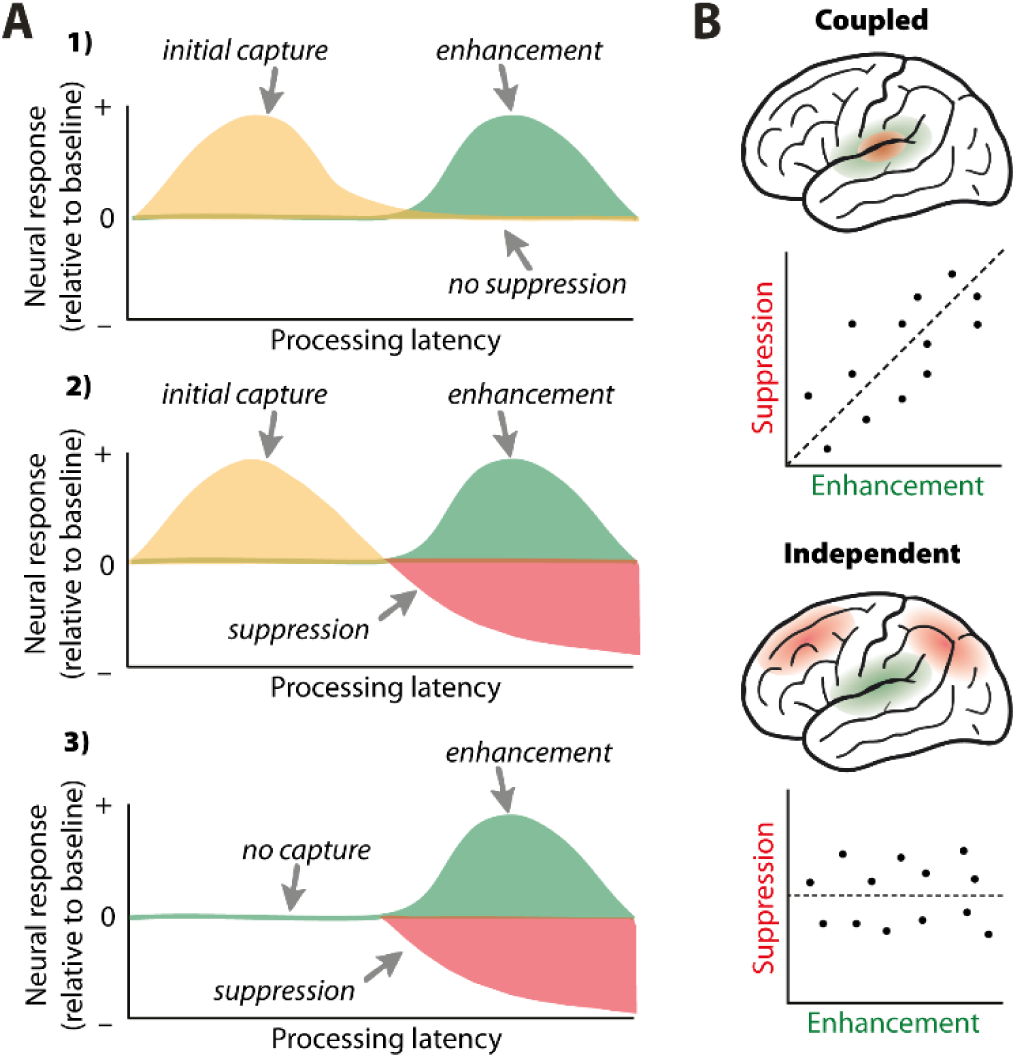
Competing theoretical predictions. **A)** Three competing models of target and distractor processing (see also (Drennan & Gaspelin, 2026). Target enhancement is highlighted in green, attentional capture by the distractor in yellow, and distractor suppression in red. **B)** Overlapping neural generators and/or correlation of enhancement and suppression would suggest coupling of the two mechanisms (top). In case of independence, no correlation and spatially separate neural generators would be expected (bottom).

Distinguishing capture from suppression is particularly challenging in auditory research. Typical paradigms employ only two competing objects (Cherry, 1953; Mandal et al., 2024; Wöstmann et al., 2016, 2017) or multiple equally salient distractors (e.g., Choi et al., 2014; Lewald et al., 2016). Such paradigms often include statistical dependencies of targets and distractors (Schneider et al., 2022) and lack neutral baselines (Wöstmann et al., 2022). Consequently, reactive auditory suppression dynamics remain largely unknown (but see Dalton & Hughes, 2014; Dalton & Lavie, 2004, 2007). This knowledge gap is critical, as everyday listening often requires selecting sounds without prior cues-a process relying on reactive filtering.

Lateralized electrophysiological responses can dissociate neural enhancement and suppression in attention. Proactive spatial attention induces differential modulations of lateralized alpha oscillatory power (∼10 Hz) related to target enhancement versus distractor suppression (Cruz et al., 2025; Wöstmann et al., 2019; Yang et al., 2024). In visual attention, the stimulus-evoked N2-posterior-contralateral (N2pc) event-related potential (ERP) component in the electroencephalogram (EEG) reflects attentional orienting (Constant et al., 2025; Eimer, 1996), and its auditory analog, the N2-anterior-contralateral (N2ac), has been associated with focusing in auditory attention (Gamble & Luck, 2011; Klatt et al., 2018). Conversely, the visual distractor positivity (P_D_) is associated with active suppression (Gaspelin et al., 2023; but see Forschack et al., 2022; Oxner et al., 2025). Some evidence from a visual attention paradigm with auditory distractors suggests that an auditory P_D_ might exist as well (Lunn et al., 2023). The neural generators of these facilitatory and suppressive responses, though investigated in animal models in a visual paradigm (Cosman et al., 2018), are largely unknown in the auditory modality.

Here, we test the mechanistic implementation of reactive auditory attention using a search paradigm with spatially separate auditory objects. The absence of foreknowledge forced participants to search for the target among distractors (Gaspelin & Luck, 2018a; Luck et al., 2021). We hypothesized that lateral targets and distractors would evoke respective neural responses associated with enhancement versus suppression (N2ac & P_D_). To contrast competing theoretical models of auditory attention (Fig. 1), we tested the relation of signatures of enhancement and suppression.

## Materials and methods

### Participants

34 participants (27 females, 32 right-handed according to self-report) took part in the study (mean age= 24 years; SD= 4; range: 19-35). Prior to the experiment, participants provided written informed consent and reported no known history of diagnosed audiological or neural deficiencies. Participants were either financially compensated at an hourly rate of€ 12 or received course credit. All procedures were approved by the local ethics committee of the University of Lübeck.

### Sample size

We performed a statistical power analysis (using Jamovi, version 2.6.26.0) to approximate sample size based on prior behavioral piloting and literature on the N2ac and P_D_ components. We inferred a smallest effect size of interest of Cohen’s *d* = 0.5. At least 34 participants were required to achieve statistical power of 0.8 with a significance level of a = 0.05 in a within-subject, two-sided test design.

### Stimuli

Auditory stimuli consisted of single-syllable spoken numbers (1-9) articulated by a female voice (250 ms duration; 44,100 Hz sampling rate; mean *fa=* 191 Hz; Fig. 2A; (Wöstmann et al., 2017). Stimuli were digitally processed using the Python libraries *slab 1.5.1* (Schönwiesner & Bialas, 2021) and *librosa 0.10.2* (McFee et al., 2015). Target stimuli were generated by multiplying the amplitude of each original waveform with a full-scale 30-Hz sinusoid at a fixed starting phase. Distractor sounds were created by shifting the pitch of the original sounds by 10 semitones using a phase vocoder algorithm *(/ibrosa.effects.shift_pitch),* resulting in a mean *fa* of 340 Hz. The final stimulus pool comprised nine original spoken numbers (controls), nine amplitude-modulated numbers (targets) and nine high-pitched numbers (distractors).

**Figure 2.**
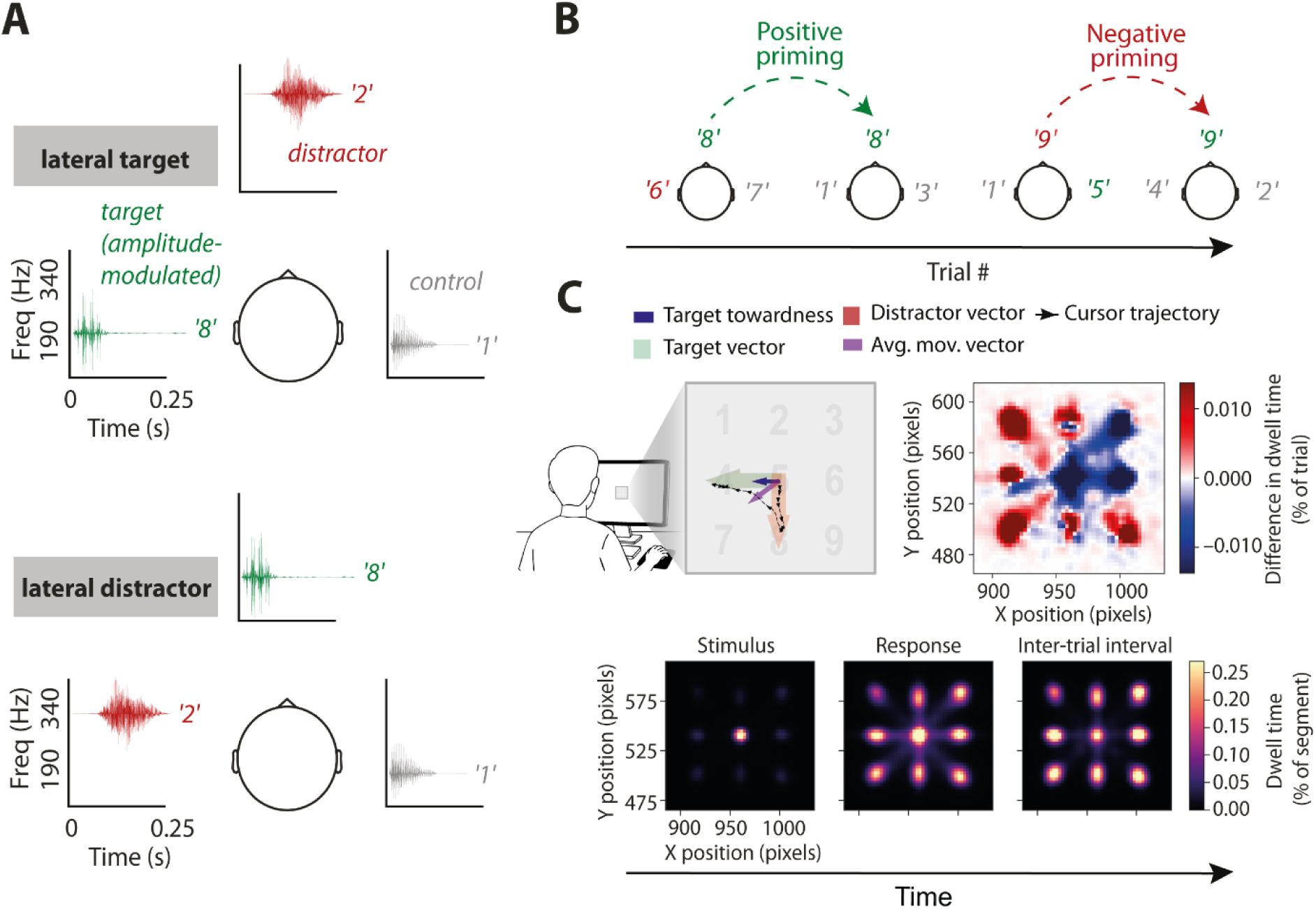
Experimental paradigm and cursor trajectory analysis. **A)** Speaker arrangement and sound characteristics. Three one-syllable numbers were presented simultaneously from three spatial positions (–90°, 0°, and +90° azimuth relative to the participant’s head). Experimental conditions included lateral target trials (top) and lateral distractor trials (bottom). **B)** Examples of target repeats (positive priming) and distractor-target switches (negative priming) within a sequence of four consecutive trials. **C)** Top left: Participants were sitting in front of a screen with a digital number pad. On every trial, the cursor trajectory used to report the target number (black trace of arrows) was averaged (purple arrow). Target towardness (blue arrow) was calculated as the scalar product of the average movement vector and the target vector (green arrow). The figure shows a trial in which the initial movement direction was aimed towards the distractor (red arrow). Top right: Difference of cursor dwell times for correct – incorrect trials. In incorrect trials, participants tended to hover around the initial cursor position in the middle of the display. Bottom: Single-trial cursor dwell times were collected and are shown here for all participants and trial phases. In this illustration, dwell times are normalized by phase duration and sampling frequency for visualization purposes.

### Auditory search task

We implemented an auditory search paradigm in which three spatially separated sounds were presented simultaneously (Fig. 2A; for a similar approach, see Orf et al., 2023). The task design resembles aspects of the well-established additional singleton paradigm in visual attention research (e.g., Bacon & Egeth, 1994; Folk et al., 1992; Gaspelin et al., 2015; Theeuwes, 1992). In this paradigm, participants search for a unique task-relevant target shape amidst several non-relevant distractors on a screen and report the orientation of a bar inside the target. In a subset of trials, one distractor (the “additional singleton”) deviates from all remaining objects in a salient but task-irrelevant dimension (e.g., color) to capture attention in a stimulus-driven way.

We transformed aspects of this paradigm into the auditory modality by having participants search for an amplitude-modulated spoken target number among irrelevant numbers and report its identity. The additional singleton was a task-irrelevant spoken number that deviated from all other sounds in frequency that was presented in 50 percent of the trials. This configuration allowed us to investigate whether a salient auditory distractor would capture attention and/or would be suppressed. Not only is this paradigm well established in visual attention research; it further allows to distinguish attentional dynamics related to capture and suppression of distractors, as well as enhancement of targets.

Stimulus lateralization within the additional singleton paradigm allows distinguishing temporally resolved neural responses to different stimulus types. It has been shown that by presenting one stimulus in front/centrally and the others lateralized, it is possible to isolate brain activity related to the lateralized stimulus at contralateral electrodes (Hickey et al., 2009; Mandal et al., 2024). Here, we employ trials with lateral target versus distractor sounds to study electrophysiological responses related to target enhancement versus distractor capture or suppression, respectively. Previous visual studies demonstrated that these conditions evoke event-related potentials which relate functionally to attentional orienting (N2ac) and suppression (P_D_), respectively (Eimer, 1996; Gamble & Luck, 2011; Gaspelin et al., 2023). By presenting less than four sounds simultaneously, the present task design allows the auditory system to effectively disentangle the pressure wave mixtures arriving at the eardrum into separate objects (Pilaszanovich et al., 2026; Zhong & Yost, 2017).

To assess enhancement and suppression behaviorally, we aimed for a fixed proportion of target repeats and distractor-target switches (further referred to as positive and negative priming trials, respectively; see Fig. 2B; Frings et al., 2015). In 20 percent of trials, the target was repeated in location and identity within two consecutive trials. In another 20 percent of trials, the distractor location and identity became the consecutive target location and identity, respectively. In the remaining 60 percent of trials, no such serial dependencies occurred (no priming trials). Due to practical constraints of generating random sequences that satisfied our criteria, the final trial sequences comprised 19.58 (SD= 0.18) percent negative priming trials, 60.83 (SD = 0.23) percent no priming trials, and 19.58 (SD= 0.23) percent positive priming trials.

The logic of this manipulation rests on selection history effects: In positive priming trials, the enhancement of target features from the preceding trial should persist, facilitating current task performance. Conversely, in negative priming trials, the suppression of distractor features from the previous trial should carry over to the present trial and reduce performance. Analyzing these history effects allows for the behavioral detection and disentanglement of enhancement and suppression mechanisms (Eben et al., 2020; Seidl et al., 2012; Tipper, 1985).

### Procedure

Participants were seated in a chair in front of a monitor (resolution: 1920xl 080; refresh rate: 60 Hz; height: 20 cm; width: 30 cm; Portable HDMI Screen, Wixamit) at a distance of ∼0.7 m. The monitor was placed below and slightly in front of the loudspeaker (15 cm) at 0° and did not obstruct the sound. Participants were instructed to keep their gaze fixated at the virtual button box on the monitor. Prior to task instructions, the EEG setup was prepared.

To ensure participants could reliably distinguish the target from the control sound, they underwent a baseline screening. Using a pseudorandomized sequence in which every number was presented at least once, 10 numbers were randomly presented across three speaker positions. Participants classified whether the heard sound was a target or a control via button press and were required to achieve 100% accuracy to proceed with the main experiment. Those who failed repeated the procedure until the criterion was met. On average, participants reached the required accuracy in 1.44 attempts (SD= 0.50).

To familiarize participants with the auditory search task, they performed 15 training trials, after which they could raise final questions about the task procedure. On every trial, participants listened to spoken numbers from three separate directions and were instructed to report the identity of the target sound via mouse click on a digital button box on the screen (Fig. 2C). The button box covered ∼2 degrees visual angle (dva; ∼2.44 cm) and the distance of the numbers relative to the center was ∼0.6 dva. In the beginning of every trial, the cursor appeared at the center of the response box (see lower panel in Fig. 2C). If a response was given later than 1.75 seconds post stimulus offset, the frame of the button box changed its color from black to red to remind the participant to respond faster. After each block, participants were allowed to take a short break, with a duration of at least one minute.

In every trial, a target sound was presented from one of three loudspeakers (Genelec: Speaker 8020D, Denmark; azimuth: –90°, 0, +90°; elevation: 0°; distance to the participants’ head: 0.85 m; Fig. 2A). Simultaneously, two irrelevant sounds were presented from the remaining two spatial locations. In 50 percent of trials, one of the irrelevant sounds was a salient distractor, which spectrally deviated from the remaining stimuli (i.e., raised in pitch by 10 semitones).

Trial-wise stimulus selection was pseudorandomized using a genetic algorithm implemented in *pygad 3.3.1,* ensuring 900 distractor-present trials, 360 positive priming trials, and 360 negative priming trials out of 1800 total trials per participant. Genetic algorithms can be used to create trial sequences that meet specific criteria (with a certain tolerance), such as balancing the distribution of priming conditions while maintaining high levels of randomization and a short run time, outperforming traditional optimization methods such as constraint satisfaction (Ihrke, 2011).

The experiment was divided into 10 blocks, each block containing 180 trials. A single trial was divided into three phases: 250 ms auditory stimulus presentation, 1750 ms response acquisition, and an average inter-trial interval of 1000 ms (randomly jittered between 750 and 1250 ms; Fig. 2C lower panel). This approach yielded an average trial duration of 3000 ms and a total experiment duration of ∼90 minutes. The experiment was implemented in *PsychoPy 2024.1.4* (Peirce et al., 2019) and conducted in a hemi-anechoic, soundproof acoustic chamber at an overall sound intensity of ∼65 dB(A).

Prior to the main task, we collected data from questionnaires (Weinstein Noise Sensitivity Scale; Speech, Spatial and Qualities of Hearing Scale), a visual Flanker task, and a passive listening task in which all sound categories (target, distractor, control) were presented sequentially in random spatial locations and order with no task relevance information. These data are not presented in the present article.

### EEG data recording and analyses

The EEG recording setup and procedure were similar to previous studies in our lab (Wöstmann et al., 2019). EEG data were recorded at 64 Ag/AgCl scalp electrodes (international 10-20 system) using Brain Vision Recorder Software (ActiChamp, Brain Products, München, Germany) and band-pass filtered online from direct current (DC) to 280 Hz against online reference channel ‘Fz’ at a sampling rate of 1000 Hz. To ensure equivalent placement of the EEG cap, the vertex electrode (Cz) was placed at 50 percent of the distance between inion and nasion and between left and right ear lobes. Impedances were kept below 20 kOhm.

EEG data were preprocessed offline using *MNE-Python 1.8.0* (Gramfort et al., 2013) and custom analysis scripts. First, we down-sampled the continuous time series data to 250 Hz and visually inspected their power density spectra. Bad sensors were manually labelled and interpolated using the spherical spline method (Perrin et al., 1989). In total, an average of 1.0 (SD: 1.28) sensors were interpolated. We then re-referenced the data to the robust average of all electrodes.

Next, non-neural components were automatically removed via independent component analysis (ICA) using */CLabe/ 0.7.0* in MNE-Python (Pion-Tonachini et al., 2019). For this purpose, a copy of the time series data was bandpass-filtered between 1 and 100 Hz and fed into an extended infomax ICA decomposition algorithm. All independent components labelled as “brain” or “other” with above 90 percent predicted probability were retained. On average, this approach resulted in a rejection of 13.26 (SD = 5.27) out of 63 ICA components. We then applied this ICA solution to the original data, after which they were bandpass filtered between 1 and 40 Hz (finite impulse response filter, FIR; zero-phase lag; hamming window).

Next, we segmented the continuous data into epochs around stimulus onset (1.0 s pre-stimulus, 1.0 s post-stimulus) and matched them with single-trial cursor trajectories. Lastly, we utilized the *autoreject* algorithm for automatic data-driven detection of bad segments (Jas et al., 2017). On average, 70.85 (SD= 74.09) epochs out of 1800 were rejected.

To isolate signatures of target enhancement from distractor capture/suppression in phase-locked neural activity (N2ac, P_D_) we split epochs into lateral target (target lateral, distractor front) and lateral distractor conditions (distractor lateral, target front; see Fig. 2A), followed by time-locked averaging across trials.

The N2ac and P_D_ have been found to show up at similar electrodes and in largely overlapping time windows (Gamble & Luck, 2011; Lunn et al., 2023). We chose electrode pair C3/4 and a time interval of 200-400 ms post-stimulus onset in our analyses for both components. Since P_D_ and N2ac are difference waves by definition, we calculated these by subtracting ipsilateral from contralateral electrode activity in the respective conditions.

### Analysis of cursor trajectory and towardness

To quantify not only how fast and accurate participants responded but how much their response was drawn to target versus distractor and control sounds, we analyzed cursor trajectories on a trial-by-trial basis (Fig. 2C). This analysis provides a continuous and direct measure of spatial attraction, complementing indirect metrics like reaction time and accuracy. Prior to this analysis, because the cursor trajectory sampling rate varied around 60 Hz, we linearly down-sampled the data to 50 Hz.

From the resampled data, we calculated an average trajectory vector for each trial by taking the mean of all x and y coordinates recorded during the response interval of each trial. Further, we defined a direction vector for the target, originating at the center of the screen and pointing to the coordinates of the target. We projected the average trajectory vector onto the target direction vector by calculating the dot product. The resulting scalar value, which is referred to as ‘towardness’ (adopted from Van Ede et al., 2020), quantifies how much the participant’s overall cursor position was aligned with the target direction vector.

Note that trials in which the target sound was a “fünf” (English: “five”) were excluded from analysis, as well as trials with implausibly large non-normalized target towardness values above 2 dva (indicating movement of the cursor way beyond the limits of the button box). We chose this value because it approximately corresponds to twice the distance between the center of the button box to its left and right boundaries. Lastly, we accounted for the variability in vector lengths in numbers positioned in the corners of the digital response pad (1, 3, 7, and 9). We normalized single-trial target towardness values by the length of their respective single-trial target vectors. Thus, a larger positive score, with a theoretically optimal value of 1.0 (perfect alignment between average cursor position and target location), indicates that the trajectory was more directed toward the target. For comparison of target towardness to conventional behavioral metrics of accuracy and response times, see Fig. S1.

### Source localization of N2ac and P_D_

We performed source localization using *MNE-Python.* A standard MRI template was used to create the head model. We defined the source space as a grid of dipoles on the white matter surface, resulting in 1026 sources per hemisphere. A three-layer Boundary Element Model was constructed with conductivities of0.3, 0.006 and 0.3 S/m for the scalp, skull, and brain compartments, respectively.

We co-registered a standard electrode montage (“easycap-M1”) to the template head model using an Iterative Closest Point algorithm based on standard fiducial locations and computed the forward solution with a minimum distance constraint of 5.0 mm from the inner skull. Consecutively, we calculated the inverse solution using dynamic Statistical Parametric Mapping. The noise covariance matrix was estimated from the pre-stimulus baseline period (–300 to –100 ms) of the epochs using Ledoit-Wolf regularization. Dipole orientation was constrained to be perpendicular to the cortical surface.

To isolate the anatomical generators of the N2ac and P_D_ components, source estimation was focused on the 200-400 ms post-stimulus time window. For each subject, the inverse operator was applied to single-trial epochs, which were then averaged to produce source estimates for lateral target and lateral distractor conditions. Similar to N2ac and Po analysis in sensor space, we calculated these components by subtracting ipsilateral from contralateral source activity in the respective lateral target and lateral distractor conditions.

Group-level statistical maps were generated by performing vertex-vise paired t-tests between contralateral and ipsilateral source amplitudes within each voxel. The resulting t-values were converted to z-scores. For visualization, statistical maps were thresholded at l*z*l > 1.96 (corresponding *top<* 0.05, two-tailed) and projected onto a partially inflated cortical surface.

### Statistical analyses

Data processing and statistical analyses were conducted using custom written Python scripts *(MNE-Python 1.8.0, scipy 1.13.0* and *statsmodels 0.14.2)* (Seabold & Perktold, 2010). Linear mixed-effects models were computed in Jamovi (The jamovi project, 2025). Prior to all analyses, trials with reaction times longer than two standard deviations from the within-subject mean were excluded (mean discarded trials = 94, SD= 19).

We evaluated the influence of distractor presence and spatial target locations on towardness using a 2×3 repeated-measures ANOVA (distractor presence, target location), reporting partial eta squared as an effect size (fl/). Follow-up post-hoc analyses and investigation into negative/positive priming effects on towardness used paired t-tests and report Cohen’s *d* as effect size. To examine the relationship between behavioral priming effects, we calculated Spearman’s rank correlations between positive and negative priming effects on towardness.

In case of non-significant results, we report the Bayes factor (BF). The BF indicates how much more likely the observed data are under the alternative (H1) compared with the null hypothesis (H0). By convention, a BF >3 begins to lend support to H1, whereas a BF< 0.33 begins to lend support to H0 (Dienes, 2014).

Statistical analysis of ERPs was done using non-parametric cluster-permutation tests (Maris & Oostenveld, 2007). For each participant, a contra-ipsilateral difference wave was calculated for both lateral target and lateral distractor conditions. These difference waves were tested against zero using a series of one-sample t-tests at each timepoint. Contiguous timepoints exceeding a threshold of *p* < 0.05 (uncorrected) were grouped into clusters. The significance of each cluster was determined by comparing its sum oft-values against a null distribution generated by 10,000 random sign-flips of the subject-level contra-ipsilateral difference.

To examine how neural activity explains trial-by-trial variability of towardness, we computed single-trial contralateral-minus-ipsilateral difference waves for each participant at electrode pair C3/4. From these single-trial difference waves, we extracted the mean amplitude within a 200-400 ms post-stimulus measurement window. N2ac/PD latency was defined as the time at which the absolute cumulated magnitude of a difference curve reached 50% of its maximum (fractional area latency). To account for the opposite polarities of these components, the negative-going N2ac difference waves were first inverted (multiplied by –1) so that both components were treated as positive-going deflections. The waveform within the 200-400 ms analysis window was then scaled to a range of O to 1 to ensure that the latency estimation is purely driven by the shape of the component rather than being biased by its peak amplitude. Single-trial N2ac latency values ranged from 0.24s – 0.37s (median= 0.30s), and their amplitudes ranged from –12.45 µV– 19.85 µV (median = –0.21 µV). Single-trial P_D_ latency values ranged from 0.24s – 0.36s (median = 0.30s), and their amplitudes ranged from –11.28 µV – 11.78 µV (median= 0.13 µV).

In two separate linear mixed-effect models, we used single-trial towardness as the response variable in lateral target and lateral distractor trials and regressed it on the following predictors: single-trial contra-ipsi ERP latency, single-trial contra-ipsi ERP amplitude, priming type (no priming, positive, negative) and absolute trial number. Additionally, the relationship between N2ac and P_D_ was assessed via Pearson correlations for both mean amplitudes and fractional area latencies. We report the standardized partial effect sizer for all relevant estimates, based on the t-value and the Sattherthwaite-approximated degrees of freedom *(df)* as: 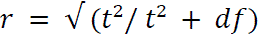 (Carlson & Furr, 2009).

## Results

In an auditory search paradigm, participants searched for an amplitude-modulated spoken target number among simultaneous distractor numbers. In 50% of trials, a high-pitched salient distractor was present. The goal of this study was to contrast competing theoretical predictions regarding the neural implementation of target enhancement versus distractor suppression, and their relation to response behavior (Fig. 1).

### Distractor presence and target position affect task performance

First, we tested whether target location (left, frontal, right) and distractor presence (present, absent) would modulate target towardness, using a 3×2 repeated-measures ANOVA. The main effect of distractor presence was significant (F_1, 33_ = 75.21, *p* < 0.001, n*_p_*^2^ = 0.70). A post-hoc paired t-test revealed that mean towardness significantly increased in distractor-present compared to distractor-absent trials (*t*_33_ = 8.53, *p* < 0.001, *d* = 0.83; Fig. 3A). While better performance in distractor-present trials might appear counter-intuitive at first glance, this finding is common also in visual search tasks and is thought to indicate successful suppression of the salient distractor, which improves target search (Gaspelin et al., 2015, 2017; Gaspelin & Luck, 2018a, 2018b; Redding & Fiebelkorn, 2024).

**Figure 3.**
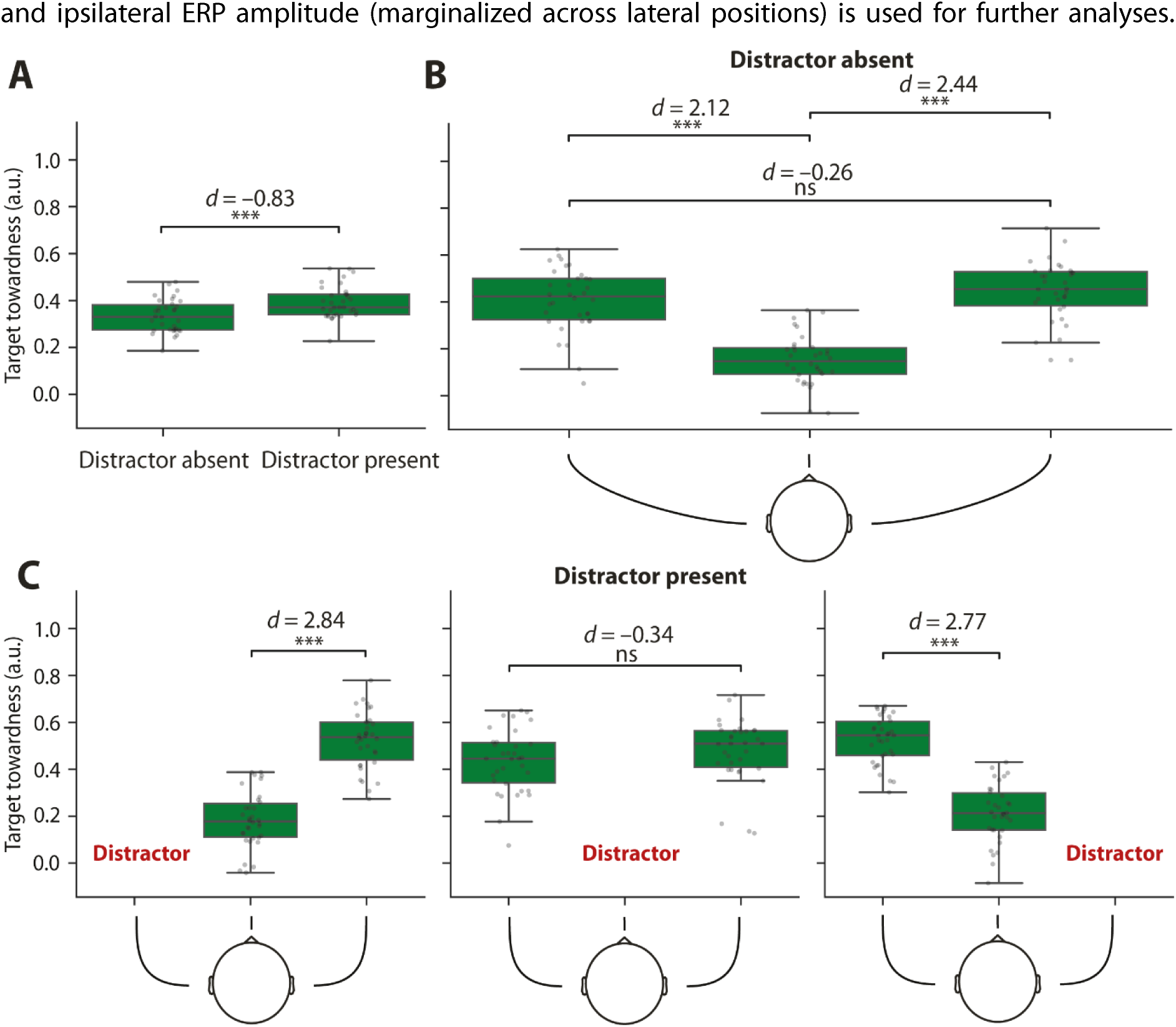
Behavioral task performance. Boxplots show towardness as a function of target and distractor location. Grey dots show mean towardness for single subjects. **A)** Distractor presence benefits towardness. **B)** In distractor-absent trials, performance decreased for frontal target locations but did not differ between left and right target locations. **C)** In distractor-present trials, a similar pattern was observed. Towardness for frontal targets decreased compared with lateral targets. Again, there was no difference in towardness for left versus right targets.* *p* < 0.05, ** *p* < 0.01, *** *p* < 0.001.

The main effect of target location was significant (*F*_2,66_ = 70.27, *p* < 0.001, n*_p_*^2^ = 0.68), whereas the interaction between target location and distractor presence was not significant (*F*_2,66_ = 1.58, *p* = 0.21, n*_p_*^2^ = 0.05). Post-hoc tests revealed that both for distractor-absent and distractor-present trials, towardness was higher for lateral than frontal targets [distractor-absent: target left vs front (*t*_33_= 10.10, *p* < 0.001, *d* = 2.12); target right vs front *(t*_33_= 10.25, *p* < 0.001, *d* = 2.44); distractor-present: target left vs front *(t*_33_= 10.91, *p* < 0.001, *d* = 2.77); target right vs front (*t*_33_ = 10.29, *p* < 0.001, *d* = 2.84)]. Target towardness did not differ for targets on the left vs right side [distractor-present (*t*_33_ = – 1.21, *p* = 0.23, *d* = –0.34, *BF=* 0.36); distractor-absent (*t*_33_ = –0.94, *p* = 0.36, *d* = –0.26, *BF=* 0.28).

Critically, these results show that our paradigm did not induce asymmetries in performance when targets/distractors were presented on the left vs right side. Thus, the contrast between contra-and ipsilateral ERP amplitude (marginalized across lateral positions) is used for further analyses.

### Neural and behavioral signatures of target enhancement and distractor suppression

We tested whether, in humans, reactive auditory attention relies on target enhancement. Behaviorally, participants’ responses were significantly more target-directed (i.e., higher towardness) in positive priming trials (i.e., repeating targets across two consecutive trials) compared to no priming trials (Fig. 4A; *t*_33_ = 3.35, *p* = 0.002, *d* = 0.42). For the EEG analysis, we focused on two distinct spatial configurations: trials where a frontal distractor was presented alongside a lateral target (upper panel in Fig. 2A), and trials where a frontal target was presented alongside a lateral distractor (lower panel in Fig. 2A). We found two time intervals during which the contra-ipsilateral difference at electrode pair C3/4 differed significantly from zero (one-sample temporal cluster permutation test; Fig. 4B). A negative difference revealed an N2ac component (0.22-0.38s, *cluster p-value* < 0.001), suggesting a relatively late enhancement of lateral targets (i.e., after obligatory ERP components; P1 & N1). Furthermore, we found a late positive difference (LPC; 0.54-0.60s, *cluster p-value* = 0.005), which has been reported for lateral targets in some previous studies (Burra et al., 2019; Gamble & Luck, 2011; Gamble & Woldorff, 2015). Source localization revealed that the neural generators of the N2ac were primarily located in the precentral, superior temporal, and superior frontal cortices (Fig. 4C). Together, these results provide evidence for reactive attentional enhancement of relevant sound and its neural basis.

**Figure 4.**
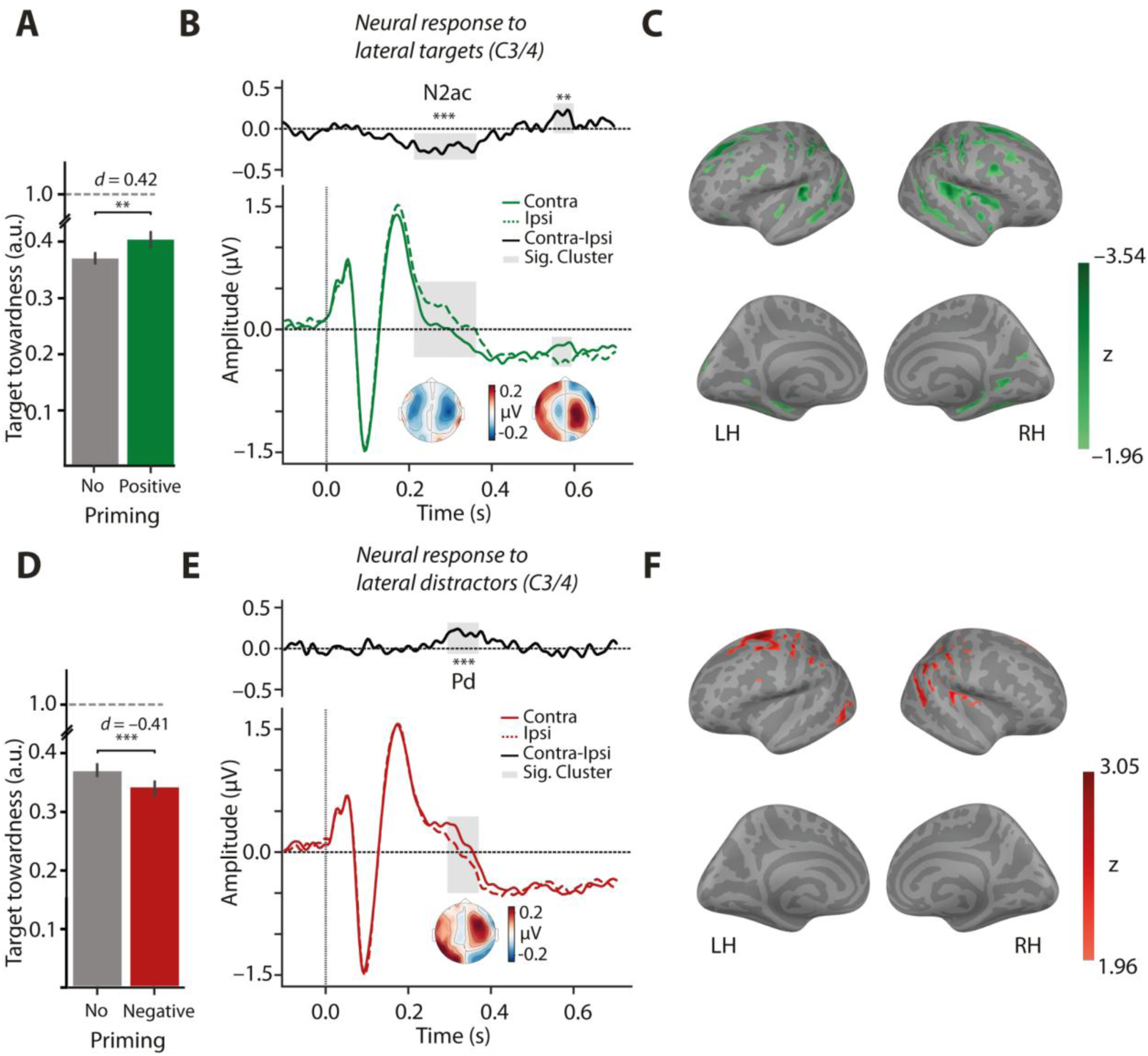
Neural and behavioral signatures of target enhancement and distractor suppression. **A)** Bars and error bars show respective average towardness ±1 SEM as a function priming. The dashed grey line represents optimal target-directed performance. **B)** Ipsi– and contralateral ERPs are indicated by dashed and solid lines (green), respectively. The black solid line in the upper panel shows the difference curve (contra-ipsilateral) between electrodes C3/4. Inset topographies show the contra-ipsilateral difference for significant clusters (highlighted in grey). **C)** Z-values for each voxel in the left (LH) and right hemisphere (RH) show the contrast of the source-reconstructed N2ac when the respective voxel is ipsi– vs contralateral to the target. Panels **D, E** and **F** follow the same principle as **A,** Band **C,** except they show behavioral distractor suppression assessed in negative priming trials **(D),** as well as neural distractor suppression in lateral distractor trials **(E&F).**

Next, we tested whether the human brain implements distractor suppression, as a complementary filter mechanism to target enhancement, during auditory search. To probe behavioral effects of distractor suppression, we utilised negative priming (i.e., distractor-target switches across two consecutive trials). Participants’ responses were significantly less target-directed (i.e., lower towardness) in negative priming trials compared to no priming trials (Fig. 4D; *t*_33_ = –5.05, *p* < 0.001, *d* = –0.41; see Fig. S2 for a control analysis including priming effects in trials with control-target switches). For the EEG analysis, we isolated trials in which the target was frontal and distractor lateralized. Analysis of the contra-ipsilateral difference revealed a significant positive cluster (Fig. 4E; 0.29-0.38s, *cluster p-value* < 0.001), that is, an auditory P_D_ component. This suggests a relatively late suppression of lateral distractors (i.e., after obligatory ERP components; P1 & N1). The neural generators of the auditory P_D_ were found mainly in precentral and superior frontal gyrus regions (Fig. 4F).

In contrast to lateralized targets, lateralized distractors evoked no LPC. Going beyond previous research that found some indications in a visual attention task that an auditory analogue of the visual P_D_ component might exist (Lunn et al., 2023), these results provide evidence that the auditory Po emerges as a robust neural signature of attentional suppression of auditory distractors.

Having established behavioral and neural indicators of selective up– and down-weighting of relevant and irrelevant sounds, we next asked whether attentional enhancement and suppression would relate across participants. First, we found no significant correlation of towardness between positive and negative priming trials, each contrasted against no priming trials *(Spearman’s rho=* 0.12, *p* = 0.50, *BF=* 0.27; Fig. S3). Second, N2ac and P_D_ latency and amplitude were not significantly correlated (latency: *Pearson’s r* = 0.32, *p* = 0.07, *BF=* 1.04; amplitude: *Pearson’s r* = –0.21, *p* = 0.24, *BF* = 0.42; Fig. S4). Third, source localization results (Fig. 4C/F) suggest largely separate neural generators of the N2ac and P_D_. These results indicate that reactive distractor suppression operates largely independently of target enhancement (for details, see Discussion).

### Neural enhancement explains trial-by-trial variation in towardness

Given the observed signatures of up– and down-weighting of respective task-relevant and task-irrelevant sounds, we asked whether these neural signatures would be mechanistically relevant for response behavior. Single-trial N2ac/P_D_ latencies and amplitudes in the time interval 0.2-0.4 s after stimulus onset were regressed on single-trial latencies and amplitudes, as well as on the absolute trial number as fixed factors and random subject intercepts.

Note that by definition, lateral target and lateral distractor trials are distinct. In total, the N2ac model included 7,243 trials with lateral targets and an average of 255 *(SD=* 21) trials per participant. The total number of trials with lateral distractors and mean per participant in the P_D_ model was 7,012 and 246 (*SD* = 22), respectively. Shorter N2ac latency predicted higher towardness (*t*_7237_ = –2.80, *p* = 0.005, *r* = 0.03; Fig. 5A). In contrast, the association of towardness with P_D_ latency was reversed in direction, but this effect did not reach statistical significance *(t*_7006_ = 1.69, *p* = 0.09, *r* = 0.02; Fig. 5B). Mean amplitudes of N2ac and P_D_ were not significantly associated with towardness (N2ac: *t*_7237_ = – 1.11, *p* = 0.27, *r* = 0.01; P_D_: *t*_7006_ = 1.04, *p* = 0.30, *r* = 0.01).

**Figure 5.**
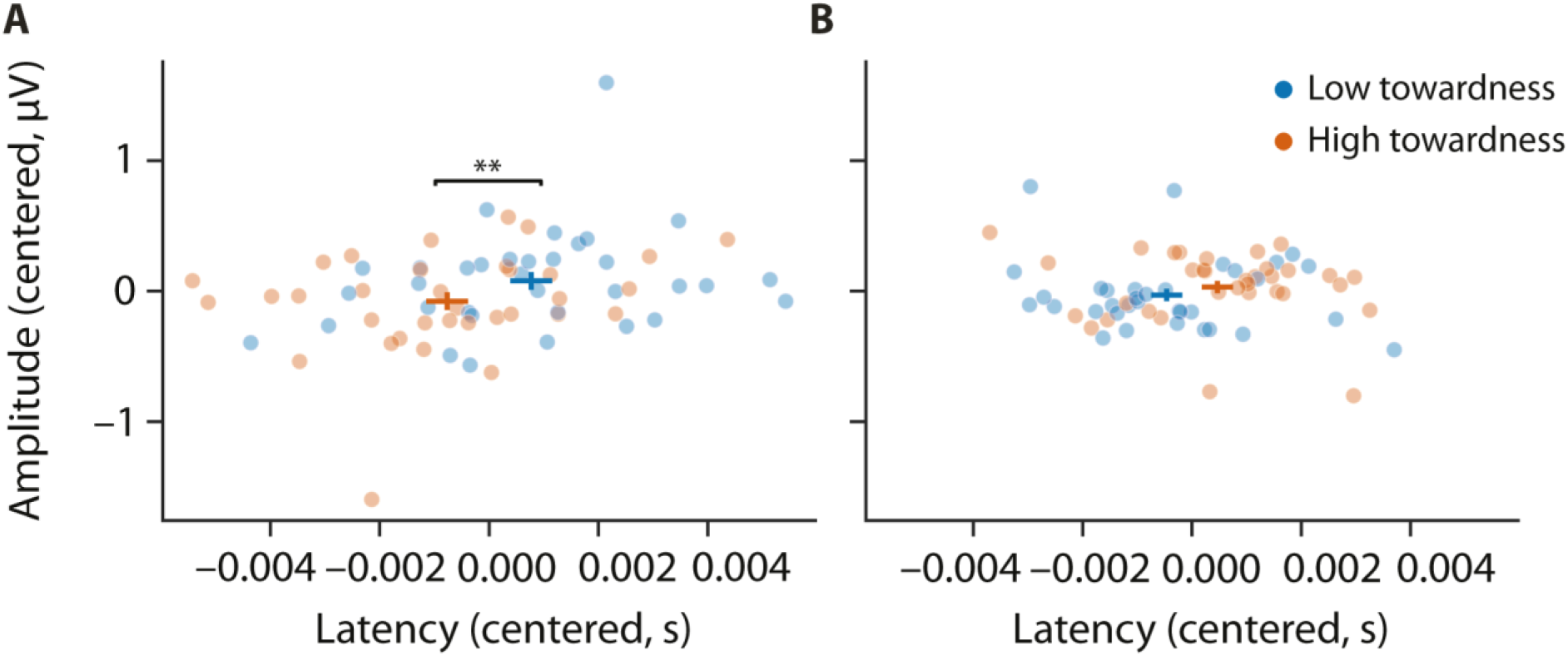
Relation of neural responses to towardness. **A)** Dots show single-subject N2ac latency (x-axis) and amplitude (y-axis). For visualization, trials were split and separately averaged for low (blue, 1^st^ quartile) and high (orange, 4^th^ quartile) towardness. Blue and orange crosses show the mean ±1 SEM of ERP amplitude and latency for low and high towardness trials, respectively. For statistical analysis, single-trial towardness was regressed on ERP amplitude and latency (see main text for details). **B)** Same as in **A),** but for P_D_ latency and amplitude. For visualization purposes, single-subject ERP amplitude and latency were zero-centered.

Since the P_D_ is associated with distractor processing but towardness mainly captures target-directed behavior, we also tested whether the P_D_ would relate to distractor-directed behavior. We regressed towardness to the distractor on single-trial P_D_ latencies and amplitudes, priming, as well as on the absolute trial number as fixed factors and random subject intercepts. Neither latency, nor amplitude was significantly associated with distractor towardness (latency: *t*_6078_ = –1.75, *p* = 0.08, *r* = 0.02; amplitude: *t*_6078_ = 0.35, *p* = 0.73, *r* = 0.004).

By definition, the N2ac is a negative deflection. To determine whether the observed effects persist when excluding trials where the contra-ipsilateral difference trends in the unexpected (positive) direction, we conducted a control analysis, using another mixed model. This analysis mirrored the initial model but was restricted to trials with a mean N2ac amplitude below 0 µV between 0.2-0.4 s. The total number of trials in this model was 3,837. Consistent with previous findings, shorter N2ac latency predicted higher towardness (*t*_3831_ = –2.55, *p* = 0.01, *r* = 0.04). Furthermore, a more negative single-trial N2ac amplitude was significantly associated with higher towardness (*t*_3831_ = –2.27, *p* = 0.02, *r* = 0.04).

## Discussion

Cognitive neuroscience has established a good understanding of how the human auditory system prioritizes relevant stimuli and suppresses irrelevant input when foreknowledge about target and distractor features is available (e.g., Lui et al., 2025; Wöstmann et al., 2019, 2021). However, the reactive neural mechanisms that resolve competition in the absence of proactive cues are not well understood. Most importantly, we here demonstrate for the first time an ERP component associated with the suppression of lateral auditory distractors during auditory attention (i.e., the auditory P_D_). Although neural distractor suppression temporally overlaps with neural target enhancement, the two processes are largely independent, emerge from distinct cortical sources, and only neural enhancement explains variability in task performance.

### The auditory P_D_ as a neural index of distractor suppression

This study is among the first to identify a contralateral positivity elicited by lateralized distractor sounds-the auditory counterpart to the visual P_D_-establishing a robust neural index of distractor suppression (Gaspelin et al., 2023). While target-related signatures like the N2ac represent established markers for auditory selection (Gamble & Luck, 2011; Klatt et al., 2018; Lewald et al., 2016; Mandal et al., 2024), evidence for a corresponding neural index of auditory suppression has remained sparse. Recent work by Lunn et al. (2023) reported a similar anterior positivity, termed the PAo, in response to lateralized sounds. In their paradigm, participants performed a central visual letter search while ignoring lateralized, task-irrelevant animal sounds. The distractors shared no features with the task-relevant visual stimuli, which is why it is not straight-forward to draw parallels to the auditory P_D_ established here. In contrast, our purely auditory paradigm utilizes distractors and targets that compete within the same modality and share fundamental acoustic (and semantic) features. This design creates the competitive environment inherent in classical selective attention paradigms and naturalistic acoustic scenes, forcing the brain to down-weight a salient yet task-irrelevant object that might otherwise be mistaken for the target. Consequently, our findings suggest that the auditory P_D_ reflects a suppressive component critical for resolving acoustic interference (Gaspelin et al., 2023; Schmid et al., 2025).

Ultimately, we found that distractor presence significantly increased towardness (Gaspelin et al., 2015, 2017; Gaspelin & Luck, 2018a, 2018b; Redding & Fiebelkorn, 2024). This behavioral benefit, coupled with the P_D_ component, suggests that the identified suppression is a functional mechanism that actively optimizes goal-directed behavior by filtering out salient distractors (although such a mechanism might also apply to non-salient distractors, see Fig. S2). Having identified an auditory P_D_, we provide a neurophysiological index to study reactive distractor processing in auditory attention research.

### Suppression of salient distractors without initial capture

A primary objective of this study was to investigate the temporal processing dynamics of auditory task-relevant versus –irrelevant objects. While attending to task-relevant targets implies signatures of enhancement in behavioral performance and neural representations (Eimer, 2014; Hamblin-Frohman & Pratt, 2026; Orf et al., 2023; Theeuwes, 2025), the evidence for distractor processing is less clear. Does a salient distractor automatically capture attention before being suppressed? Current opposing theoretical frameworks of attention, largely derived from the visual attention literature, predict contrasting outcomes (Drennan & Gaspelin, 2026).

One model suggests that salient distractors initially capture attention, followed by rapid disengagement (see first model in Fig. 1A; Theeuwes, 2010). Neurally, this should manifest in either an early enhancement of sensory-evoked responses (N1/P1 component; Berti, 2013; Chait et al., 201O; Naataanen, 2018), or an early distractor-related N2ac (analogous to Forschack et al., 2022a; Liesefeld et al., 2017; McDonald et al., 2013; Van Moorselaar & Slagter, 2019). An alternative model assumes active suppression occurs only after initial capture (see second model in Fig. 1A; Moher & Egeth, 2012). Neurally, suppression would be reflected in larger contra-than-ipsilateral ERP amplitude (i.e., a P_D_ component), reflecting active termination of distractor processing (Gaspelin et al., 2023). Both models share the assumption that salient distractors inevitably capture initial resources and that suppression is, if observed at all, a reactive process (Beck & Hollingworth, 2015; Gaspelin & Luck, 2019; Hickey et al., 2006).

Ultimately, a third model posits that salient distractors can be suppressed without initial capture (see third model in Fig. 1A; Gaspelin et al., 2025; Sawaki & Luck, 2010), which stands in agreement with our results. We found no early sensory modulation of the N1/P1 components by lateral distractors, nor an initial N2ac preceding suppression. Instead, the earliest neural signature of distractor processing was the P_D_ component, which emerged relatively late (200-400 ms) and in a nearly identical time window as the target-related N2ac. The simultaneous emergence of enhancement and suppression suggests that the auditory system bypasses the serial process of initial capture and subsequent suppression. Instead, our findings indicate that as soon as the acoustic scene is parsed, the human brain down-weights the distractor’s neural representation while it engages the focus of attention. This distinction aligns with a dual-mode framework proposed by Mandal et al. (2024), which posits that auditory attention can operate through either initial capture or direct suppression; our findings specifically provide evidence for the latter mechanism.

The neural pattern of suppression without capture is mirrored in the behavioral data: distractor presence increased towardness-a finding in line with previous studies reporting benefit of distractor presence (Gaspelin et al., 2015, 2017; Gaspelin & Luck, 2018a, 2018b; Redding & Fiebelkorn, 2024). While this benefit may partially stem from low-level spectral release from masking (Brungart, 2001; Miller, 1947)-where a spectrally more distinct distractor reduces energetic interference at the basilar membrane-the behavioral profile reinforces the neural evidence that the distractor’s attentional priority was successfully reduced. Recent work further strengthens the P_D_ as a marker of such suppression; for instance, Schmid et al. (2025) demonstrated that early broadband high-frequency activity triggered by a distractor predicts the strength of the subsequent P_D_.

However, alternative interpretations of the P_D_ exist. It might reflect feature disambiguation due to local contrast rather than active suppression (Forschack et al., 2022a). Furthermore, the interpretation of the P_D_ as a signature of suppression rests on the assumption that the control stimulus serves as a neutral baseline. In many search paradigms, target and control stimuli share most features (see Oxner et al., 2023) for a view on target feature enhancement), meaning that enhancement of the control stimulus could make the distractor appear suppressed by comparison (Kerzel & Burra, 2020; Kerzel & Huynh Cong, 2023). Establishing a truly neutral baseline remains a persistent challenge in separating these mechanisms (Wöstmann et al., 2022). While our data show no initial capture, such a process may still exist, for instance in the very first instance of distractor occurrence (e.g., the first trial) before rapidly evolving into suppression (Zhang & Gaspelin, 2026).

### Independence of auditory enhancement and suppression

Using a spatial loudspeaker setup that allows us to distinguish lateralized neural responses to targets and distractors, we here provide evidence that in auditory reactive attention, enhancement and suppression are implemented largely independently: Both behavioral and neural signatures of target enhancement and distractor suppression-indexed by positive and negative priming, as well as N2ac and P_D_, respectively-were unrelated at the between-subject level. Critically, the N2ac sources were primarily localized to superior temporal and superior frontal cortices, while the P_D_ seemed more strongly generated in superior frontal and precentral brain regions. This spatial divergence suggests that while target selection primarily involves sensory processing regions, distractor suppression recruits frontal cognitive control regions, emphasizing that inhibition in attention is not a purely sensory phenomenon but a higher-order control process (see also *Geng, 2014; Hasegawa et al., 2004; Shimamura, 2000; Suzuki & Gottlieb, 2013)*. Lastly, only N2ac, but not P_D_ latency, predicted behavioral performance. This agrees with a previous auditory attention study where only the continuous neural tracking of target speech explained performance (Orf et al., 2023). Our results are consistent with recent evidence that behavioral signatures of target enhancement and distractor suppression vary independently across individuals, which challenges shared-mechanism accounts (Khodayari et al., 2026).

Empirical evidence suggests that neural markers of suppression operate independently of target selection and support the existence of functionally distinct neural populations dedicated to these two mechanisms. For instance, Wöstmann et al. (2019) utilized EEG to demonstrate that lateralized auditory distractors elicit reversed alpha lateralization compared to lateralized targets. This mechanistic divergence is further supported by Yang et al. (2024), who identified distinct alpha-mediated processes underlying target enhancement and distractor suppression during spatial attention deployment. Likewise, Seidl et al. (2012) used functional magnetic resonance imaging to show that enhancement and suppression both contribute to selective visual attention. Contrary to the assumption that distractor suppression is contingent on target selection (Noonan et al., 2018), our findings suggest that they are functionally and anatomically distinct, yet complementary mechanisms, both serving the aim to optimize goal-directed behavior.

## Conclusion

While current theories of selective attention emphasize the necessity of filtering irrelevant sensory inputs in absence of proactive cues, the reactive neural mechanisms that resolve competition in the auditory system have remained largely unclear. Employing a paradigm that dissociates stimulus-driven enhancement from distractor suppression, we demonstrate that two independent neural signatures-the N2ac and the newly established auditory P_D_-characterize the brain’s reactive response during selective listening. These components emerge within largely overlapping time intervals without evidence of prior attentional capture, with suppression recruiting distinct cortical generators. Collectively, these results support models of auditory attention where reactive enhancement and active suppression function as parallel, independent neurocognitive operations.

## Data accessibility

All data are available from the corresponding authors upon request and will be made available on osf.io upon publication.

## Acknowledgements

This work was supported by Deutsche Forschungsgemeinschaft (grant number WO 2371/1-2, to M.W.)

## Supplemental Materials

**Figure S1.**
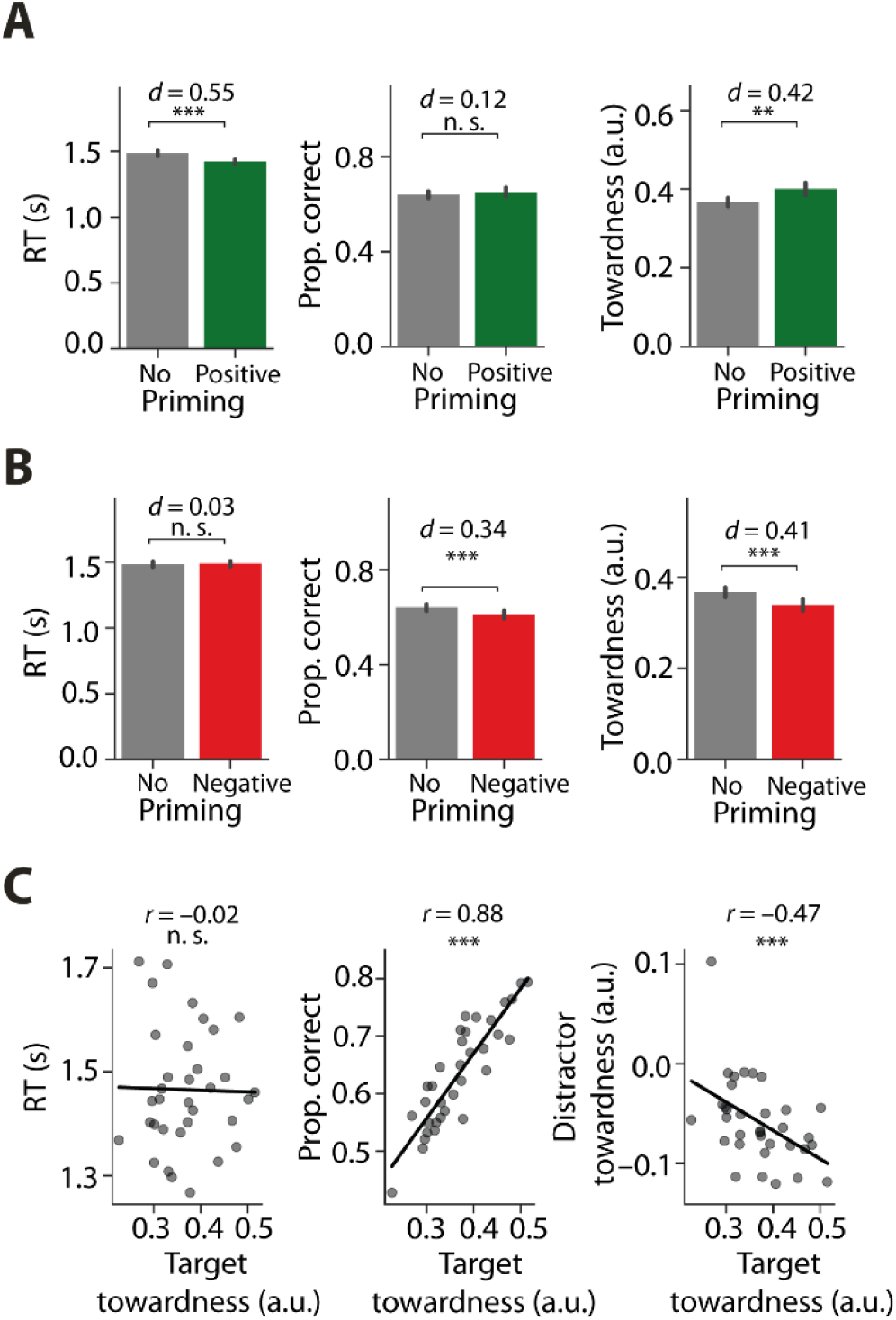
Comparison of target towardness to reaction time and accuracy. **A)** Comparison between no priming (grey) and positive priming (green) conditions across three behavioral measures: response time (left), accuracy (middle), and target towardness (right). Bars and error bars indicate mean and ±1 SEM, respectively. Significant differences between paired conditions are denoted by asterisks (*** p < 0.001, ** p < 0.01, * p < 0.05, n.s. p > 0.05), and effect sizes are reported as Cohen’s *d.* **B)** Same as A but for negative priming. **C)** Scatter plots illustrate the relationship between mean target towardness and mean response time (left), accuracy (middle), and distractor towardness (right). Each data point represents an individual subject’s mean. Black lines indicate ordinary least squares regression lines. Pearson correlation coefficients rand corresponding p-values are provided for each comparison.

**Figure S2.**
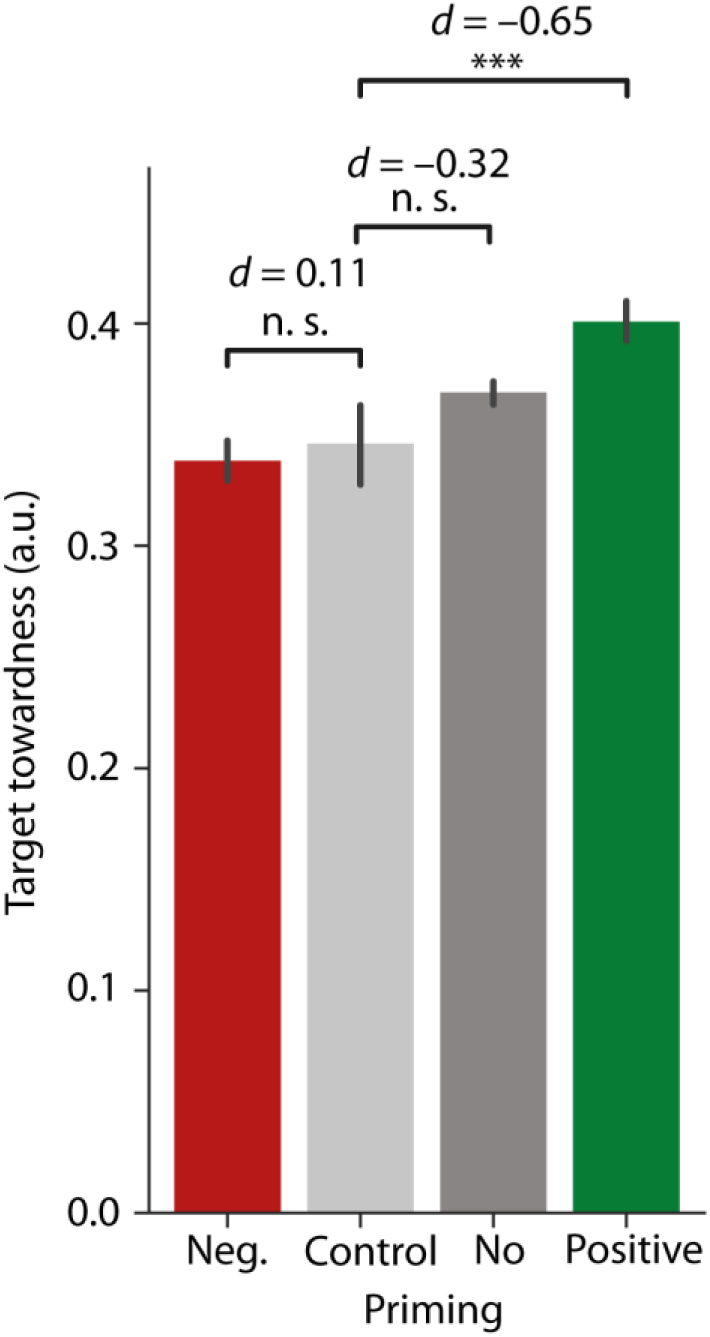
Priming effects for control sounds (i.e., non-salient task-irrelevant stimuli). Mean target towardness is shown across four priming conditions. The control priming condition specifically represents trials where the current target matches a control sound from the preceding trial in location and identity. Note that the number of trials that underlie the control priming category is one third of the size of negative and positive priming trials, and that statistical interpretability is therefore limited. Error bars represent ±1 SEM. Statistical significance between conditions was evaluated using paired-samples t-tests, with p-values adjusted for multiple comparisons using Bonferroni correction. Only the comparisons between control priming against negative, no, and positive priming are visualized. Effect sizes are reported as Cohen’s *d.* Significance levels are denoted as follows:* *p* < 0.05, ** *p* < 0.01, *** *p* < 0.001, n.s. *p* > 0.05.

**Figure S3.**
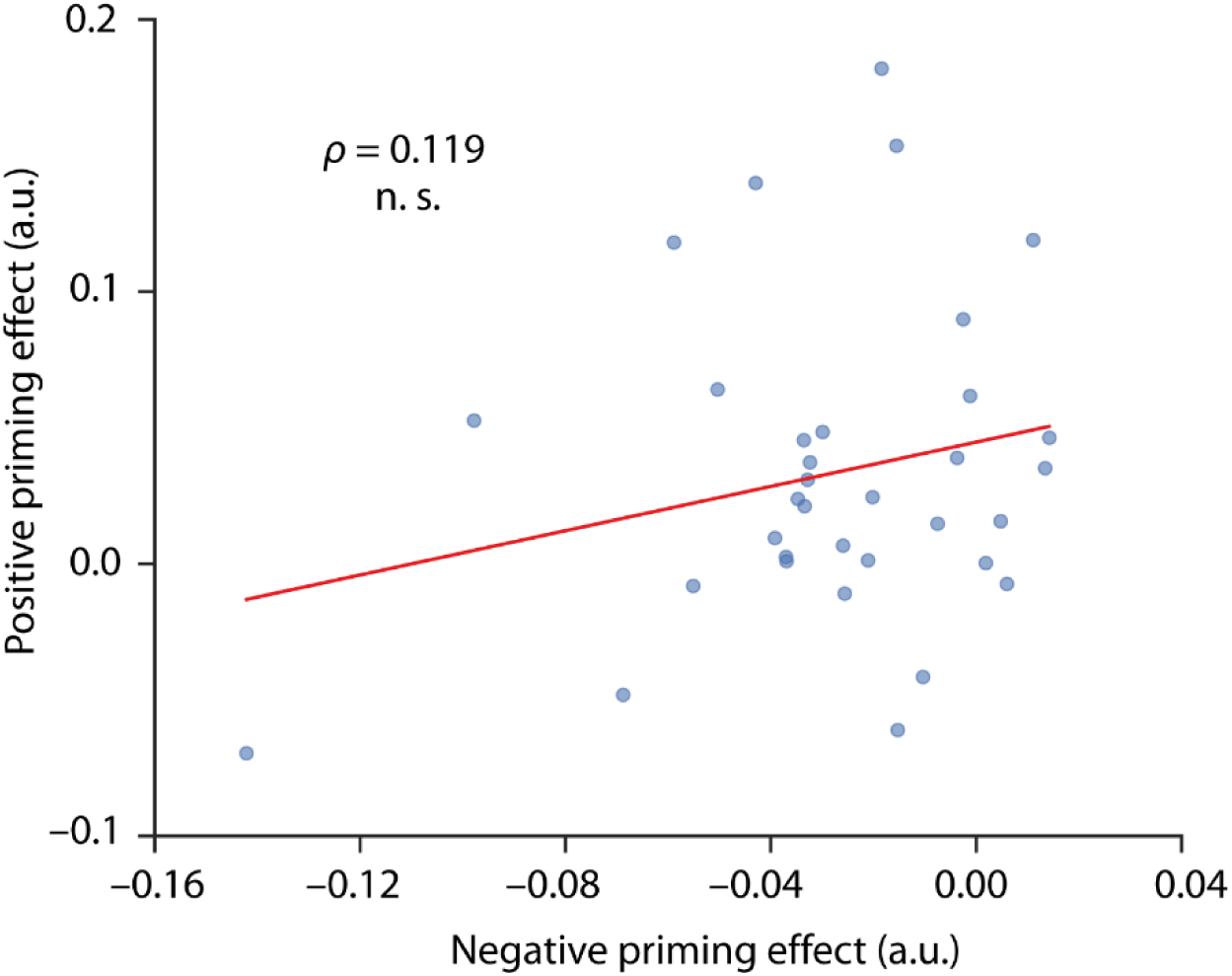
Correlation between negative and positive priming effects across participants. Each data point represents a single participant. The positive priming effect is defined as the difference in towardness between the positive priming and the no priming condition. Similarly, the negative priming effect is defined as the difference in target towardness between the negative priming and the no priming condition. The ordinary least squares regression line is shown in red. The relationship was tested statistically using Spearman’s rank correlation. n.s. *p* > 0.05.

**Figure S4.**
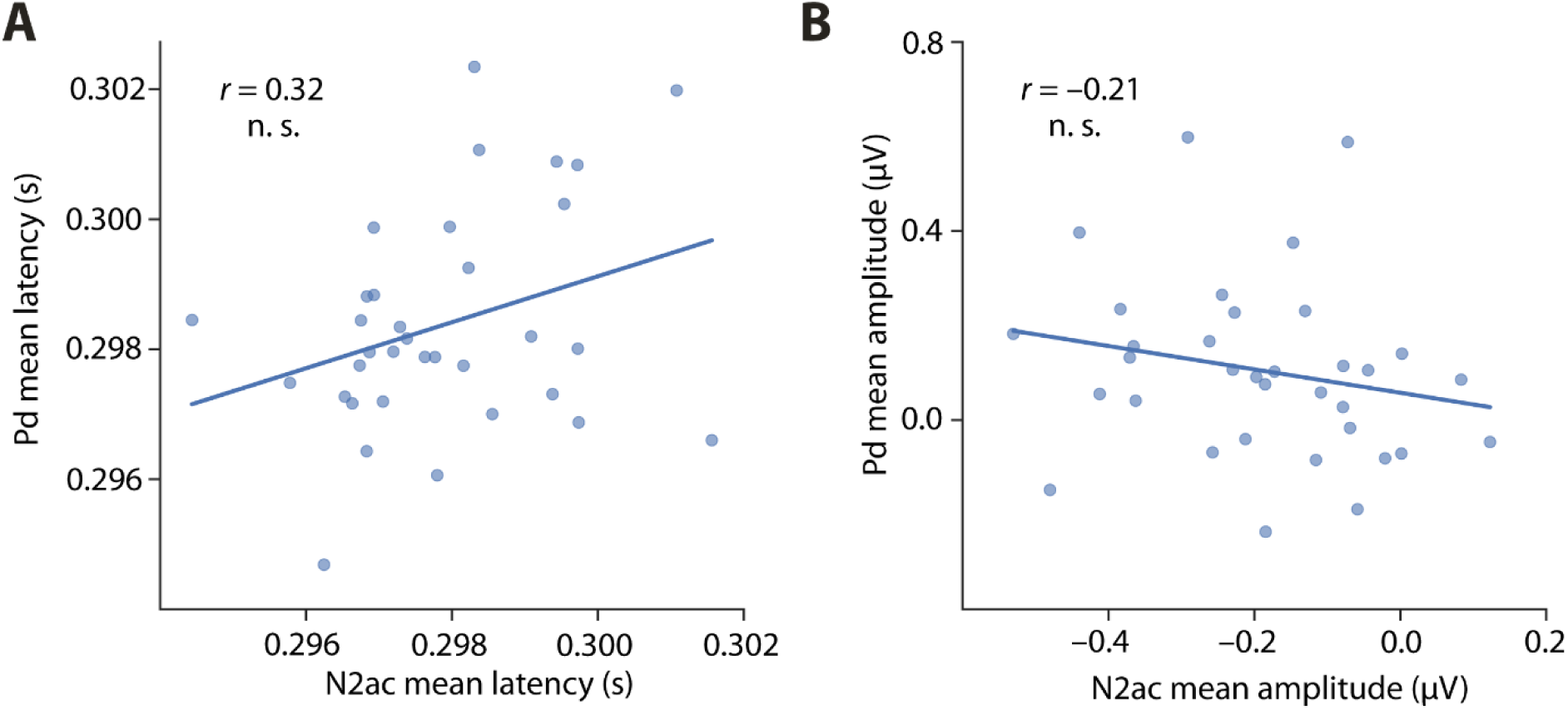
Correlation of Po and N2ac latency and amplitude across participants. Scatter plots illustrate the relation between individual N2ac and Po metrics. Each point represents a single participant. **A)** The relationship for single-trial mean latency, with the N2ac mean latency on the x-axis and Po mean latency on the y-axis. **B)** The relationship for single-trial mean amplitude, with the N2ac mean amplitude on the x-axis and Po mean amplitude on the y-axis. The ordinary least squares regression line is shown in blue. The relationship was assessed using Pearson’s correlation. n.s. *p* > 0.05.

**Figure S5.**
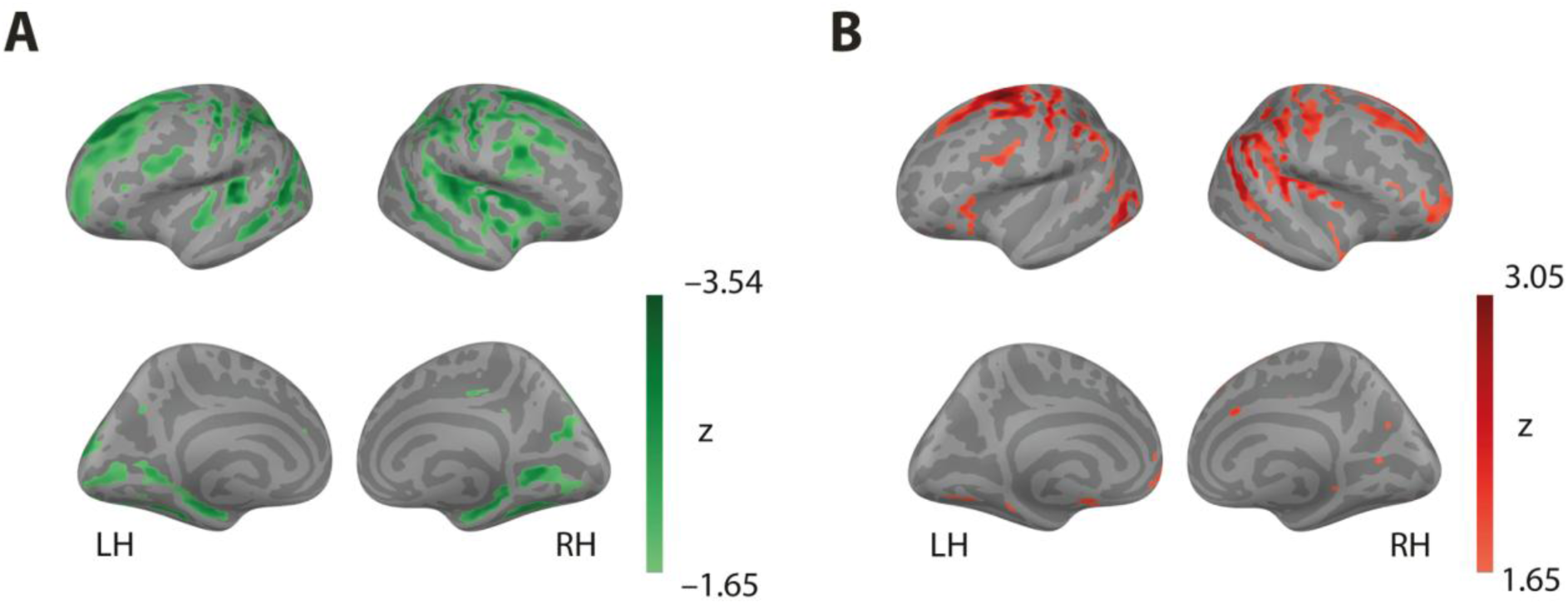
N2ac and Po source estimates. Similar to Fig. 4C/F, but for visualization, the statistical z-maps were thresholded at absolute values of l*z*l > 1.65 (corresponding *top* < 0.05, one-tailed) and projected onto a partially inflated cortical surface. Source estimates were averaged between 200 – 400 ms post-stimulus onset. **A)** Z-values for each voxel in the left (LH) and right hemisphere (RH) show the contrast of the source-reconstructed N2ac when the respective voxel is ipsi-vs contralateral to the target. **B)** Z-values for each voxel in LH and RH show the contrast of the source-reconstructed Po when the respective voxel is ipsi– vs contralateral to the distractor.

